# Immune priming can prevent WNV establishment in *Culex quinquefasciatus* mosquitoes: evidence for immune priming based reversal of WNV-mediated immune suppression

**DOI:** 10.1101/2021.05.05.442826

**Authors:** Marcus Blagrove, Seth M Barribeau

**Affiliations:** Department of Ecology, Evolution, and Behaviour, Institute of Infection, Ecology, and Veterinary Studies, University of Liverpool, Liverpool, UK

## Abstract

Mosquito-borne infectious diseases cause wide-spread loss of life and livelihood often in low-income settings. However, control of mosquito-vectored viral diseases such as West Nile virus (WNV) and Japanese encephalitis virus (JEV) remains challenging. Here we use an existing feature of the insect immune system to effectively vaccinate *Culex quinquefasciatus* mosquitoes against WNV infection. We find that priming mosquitoes by exposure to inactivated WNV reduces their likelihood of developing transmissible infections of WNV after live infection. We used RNA sequencing to identify gene expression in response to WNV and JEV infection, and the role of prior priming exposure on constitutive and induced expression on infection. Infection with either Flavivirus causes broad suppression of gene expression. WNV and JEV infection resulted in suppression of different suites of genes with notable immune genes, such as antimicrobial peptides, being strongly suppressed on WNV infection. We hypothesise that the increased resistance to WNV infection seen in primed mosquitoes may be the result of priming nullifying the immune suppression found in non-primed WNV-fed mosquitoes, potentially through greater expression of mRNA regulatory genes such as cap-binding proteins in primed mosquitoes.

**Author summary:** Mosquitoes vector many devastating infectious diseases. Two such vectored viral diseases are West Nile virus (WNV) and Japanese encephalitis virus (JEV). Control of these diseases remains challenging, and no vaccine exists for WNV. Here, we tested whether we could instead vaccinate the mosquitoes against WNV. By injecting mosquitoes with dead WNV we found that we could reduce the number of infected mosquitoes by half. We then used whole-genome RNA sequencing to identify which genes are transcribed, which will help us understand genes that are important for this form of insect immune priming, and for responses to normal WNV and JEV infection. We found that WNV suppresses the expression of many immune genes but these genes are expressed normally in vaccinated mosquitoes. Our findings expand our understanding of mosquito infection with these viruses but also demonstrate how prior exposure to a disease can produce lasting protection.

## Introduction

Mosquito-borne viral diseases infect more than 135 million people every year[1], causing the loss of 2.7 million years of human life *per annum*. Additionally, arboviruses cause significant economic loss by harming livestock and ecological consequences when infecting wildlife. These effects are compounded by the recent and continual expansion in geographic range of most arboviruses, such as Zika virus [2] and the zoonotic West Nile virus [3].

*Culex quinquefasciatus* is an especially important vector of both WNV[4] and JEV[5], as it is both a highly competent vector for both viruses and is a generalist; feeding on birds, humans, and non-human mammals [6]. *Cx. quinquefasciatus* therefore contributes significantly to both the reservoir maintenance and as a bridge-vector to humans and other dead-end mammals.

Control of mosquito-borne viruses has proven problematic. Specific treatments and vaccines are only available for very few arboviruses; consequently, most control strategies rely on vector-killing such as larval rearing site destruction and adult killing [7]. Recently however, there has been an increase in research focused on reducing the vectorial capacity of a population by e.g. heritable bacterial infections which suppress viral development [8,9], and other anti-viral gene drive strategies [10]. These methodologies recognise the difficulty in eradicating vectors as well as the potential ecological consequences of exterminating vectors.

Here, we explore another route to viral control within the vector, manipulation of the mosquito immune system. Among the most enigmatic features of insect immunity is the ability of some species to produce a lasting and indeed specific form of immune memory, often termedtermed immune priming [11]. Immune priming in insects, and other invertebrates, is analogous to vertebrate immune memory, producing greater protection to a pathogen on secondary exposure than on an initial exposure. This benefit can be long lasting, and can be transferred from parents to offspring [12]. Immune priming has been described in *Anophaline* mosquitoes to *Plasmodium* [13,14], and *Aedes aegypti* to Dengue virus [15]. While not all species show this priming response, the diversity of invertebrates with some form of immune priming is striking [11].

Immune priming can confer varying degree of protection. In the red flour beetle *Tribolium castaneum* immune priming produces specific protection down to the level of the strain of *Bacillus thuringiensis* [16]. This specificity comes with an associated cost: immune priming to one pathogen can result in greater susceptibility to other pathogens [17]. The nature of priming specificity and potential costs in greater susceptibility is especially important when exploring immune priming in vector species. As mosquitoes can vector multiple pathogens and parasites there is the potential for multiplicative benefit through cross-protection to other pathogens, or conversely, to replace any benefit if priming against one pathogen results with greater competence to other pathogens.

West Nile virus (WNV) and Japanese encephalitis virus (JEV) are mosquito vectored flaviviruses that both primarily infect non-human vertebrates such as birds and pigs. While these flaviviruses rarely cause symptomatic disease in humans, when they do, they can be highly virulent. JEV infects 45,000 people a year, killing 10,000 of them[18]. These values are thought to be underreported [19–21]. WNV is broadly distributed and spread to North America in the late 1990s where it caused the largest outbreak in 2002 with 5.9% case mortality (reviewed in [22,23]). Both JEV and WNV can cause long-lasting neurological sequelae [19,24].

Immune priming appears in diverse species, but the mechanisms that underpin it remain largely unknown. Here, we explore the transcriptomic responses of mosquitoes primed against the vectored human disease West Nile Virus, and their subsequent response to an infectious bloodmeal containing WNV or a related Flavivirus, Japanese Encephalitis Virus (JEV).

## Materials and methods

### Mosquito rearing and incubation

*Culex quinquefasciatus* (Muheza strain, G37) eggs were hatched in deoxygenated water and fed with brewer’s yeast tablets. They were allowed to emerge and mate in 30 x 30 x 30 cm BugDorm cages in University of Liverpool and incubated at 27°C, 12:12 light:dark and 70% RH. After approximately seven days post emergence, where they were transported to the CL3 infection insectary (Liverpool School of Tropical Medicine) and incubated at 27°C, 12:12 light:dark and 70% RH, where they remained until sacrifice.

### Virus strains and culture

Japanese encephalitis virus (JEV, Muar strain), was cultured in Vero cells maintained in Dulbecco’s modified Eagle’s minimal essential medium (DMEM) (Sigma-AldrichCorp., St Louis, MO, U.S.A.) containing 10% heat-inactivated fetal calf serum (FCS), 2 mm L-glutamine and 50 μg/mL penicillin/streptomycin. West Nile virus (WNV, NY-99 strain) was cultured at Public Health England, Porton Down, Surrey, in Vero cells.

### Priming treatment

At seven days post emergence, in the infection insectary female adults were removed, separated into two equal groups and injected intrathoracically using a Nanoject (Drummond, USA) with heat-deactivated WNV or control (sham) preparations containing Dulbecco’s Modified Eagle Medium (Thermo Fisher Scientific, USA). All injection volumes were 69nL, and WNV was diluted to a titre of 1×10^8^ PFU/mL using Dulbecco’s Modified Eagle Medium. Each mosquito was thus injected with approx. 6,900 virions. Immediately prior to aspiration, both preparations were heat-treated at 60°C for 30 minutes to inactivate the virus[25]. Once injected, females were transferred and counted into 1L cylindrical polypropylene DISPO-SAFE containers (Microbiological Supply Company, UK), with a fine mesh covering the container opening, and provided sucrose *ad libitum*.

### Competence experiment

At seven days post-injection (14 days post emergence), female adults which had undergone the above priming treatment and had been incubated for 24 h with no access to sugar, were blood-fed with WNV. Blood meals (heparinised human blood, NHS transfusion service, Speke) containing WNV at 1×10^6^ PFU/mL were provided for 3 hours using a feeding membrane odorised by rubbing against human skin. Unfed adults were removed from the cage, and the fed mosquitoes were incubated at 27 °C and 70 % RH for seven days.

At seven day post-feeding (21 days post emergence), mosquitoes were anaesthetised with FlyNap (Carolina Biological Supply Company, Burlington, North Carolina, USA), and their saliva was extracted by inserting their proboscis into a capillary tube containing mineral oil. Expectorate was collected from a total of 78 heat-deactivated WNV injected females and 76 sham injected females. RNA was extracted from the expectorate using TRIzol reagent (Thermo Fisher Scientific). cDNA was generated using Superscript Vilo (Thermo Fisher Scientific).

Taqman (Thermo Fisher Scientific) quantitative reverse transcription polymerase chain reaction (qRT-PCR) was used to detect the presence of viral RNA in the samples. Primer and probe sets were as follows: WNV, sense: 5′-CCA CCG GAA GTT GAG TAG ACG-3′, anti-sense: 5′-TTT GGT CAC CCA GTC CTC CT-3′, probe: Cy5-TGC TGC CTG CGG CTC AAC CC-BBQ. Cycling regimen: 1 min at 95 °C followed by 40 cycles of 95 °C for 5 s and 60 °C for 8 s [26].

### RNA sequencing experiment

At seven days post injection (14 days post emergence), a separate batch of adult female mosquitoes (which had also undergone the above priming treatment) were provided with one of three blood meals: 1) Blood only; 2) Blood containing WNV at 1×10^6^ PFU/mL; or 3) Blood containing JEV at 1×10^6^ PFU/mL. This resulted in six conditions (two priming conditions × three blood meal conditions, summarized in table 1). The blood used was heparinized human blood obtained from NHS transfusion service in Speke, Merseyside, UK. Blood meals were provided for three hours using a Hemotek feeding system and ‘Hemotek feeding membrane’ (Hemotek Ltd, UK) odorised by rubbing against human skin (approximately 0.3 and 0.4 log decrease in viral titre for WNV and JEV respectively was observed over this period, measured by qRT-PCR). Unfed females were removed from the cage.

**Table 1.**
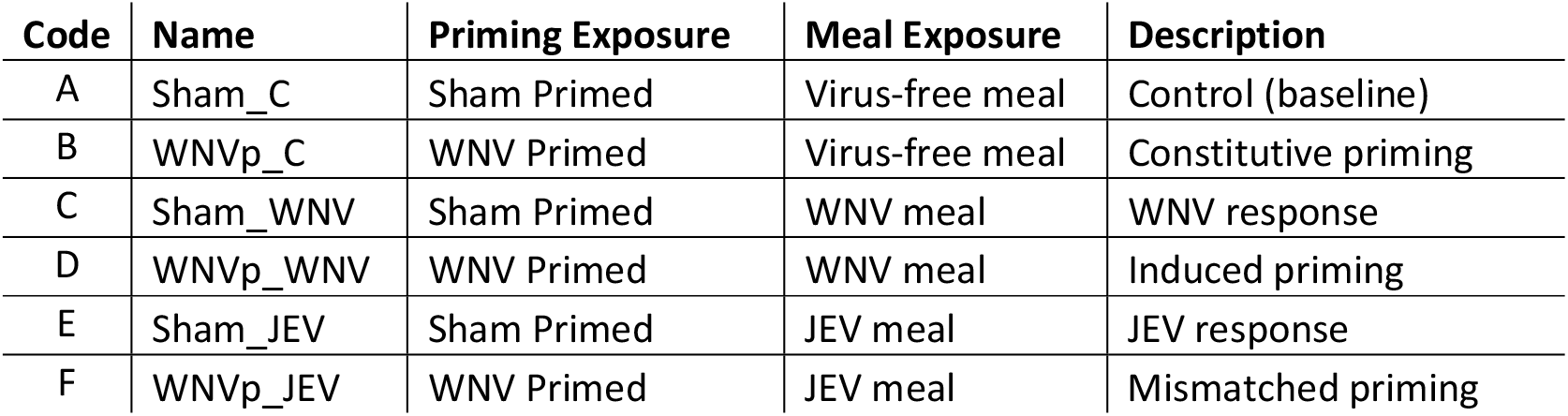
Summary of treatment conditions.

Two days after blood feeding (16 days post emergence), for each of the six conditions listed in table 1, 20 mosquitoes were anesthetized with FlyNap (Carolina Biological Supply Company, USA) and sacrificed into 10 pools of two (= total of 60 pools of two). RNA from these pools was extracted using Zymo Quick-DNA/RNA Pathogen kit (Zymo Research, USA) as per manufacturer’s instructions. RNA sequencing used the SMART-Seq v4 Ultra Low Input RNA Kit (Clontech) by Novogene (Cambridge, UK). Sequencing used the Illumina HiSeq platform.

### Bioinformatics

We trimmed the raw reads using TrimGalore with default settings and mapped the resulting trimmed reads to the *C. quinquefasciatus* genome (CpipJ2.47) using STAR (version 2.7.1a) [27] in GeneCounts mode with the associated annotation gff file. Resultant counts tables per library were combined into a single table and we used edgeR [28,29] (v 3.30.3) to identify differentially expressed genes. Our design allows for multiple different exactTest comparisons in edgeR that address different elements of our question. We used a FDR cutoff of 0.05 for tests of significance.

#### Comparisons

1. As a baseline, we compared group A (unprimed and fed a virus free meal) against all other groups.
2. We further compared expression of individuals primed against WNV that fed on a WNV infectious meal (D) to those that were not primed but also received an infectious WNV meal (C) to compare the normal WNV response to that produced in primed individuals (C vs D).
3. We contrasted individuals that were primed but not infected (B) to those that were both primed and exposed to WNV (D) to identify how secondary exposure differed to constitutive priming.
4. We explored the consequences of mismatch between priming exposure and infectious dose in two contrasts. We first compared those that were not primed and fed infectious dose of JEV (E) to those that were primed against WNV but infected with JEV (F). Second, we compared priming match (D) to mismatch (F).

We took gene ontology terms from VectorBase and used topGO (v. 2.40.0 [30]) to identify enriched ontology terms. We calculated weighted, classic, and parent-child test statistics for each GO term, but only retained those that were significant under the more stringent parent-child Fisher’s exact test result. We additionally examined the network structure and co-expression features of the whole dataset using CEMiTool [31,32] with protein interactions retrieved from String-DB [33] and annotations from gProfiler [34] with vst adjustment for mean-variance correlation, and used Spearman rank correlations to cluster gene expression. To clarify which genes are differentially expressed and the gene ontology terms are shared among comparisons we used upset plots (UpSetR v1.4.0 [35]). We used R (v 4.0.2 [36]) for all statistical analyses. In order to further contextualize our findings, we compared the differentially expressed genes and overrepresented gene ontology terms from our study to those in Bartholomay *et al*. [37], Shin *et al*. [38], and Pradakar *et al*. [39].

Finally, to understand whether immune activation by direct exposure to virus or by priming broadly affects microbes we used idSeq [40]. We uploaded all reads that did not map to the reference genome to the idSeq servers and downloaded the resulting tables that contain the total number of reads that map to non-Eukaryote sequences in the NCBI nucleotide or peptide databases. The number of reads to the nr and nt databases were summed over each taxa and then analysed using classic measures of ecological diversity (Shannon’s Index, species richness) using the R package vegan [41]. Here we are treating the number of reads mapping to a taxa as the count of that taxa in that sample. As species identifications are not always possible, we calculated diversity metrics at genus level.

## Results

Immune priming by injecting adult females with heat inactivated WNV suppressed live WNV infection. We found that exposure to a heat inactivated WNV injection resulted in approximately half the number of mosquitoes with detectable WNV in their saliva (Fig 1) than sham primed mosquitoes.

**Fig 1.**
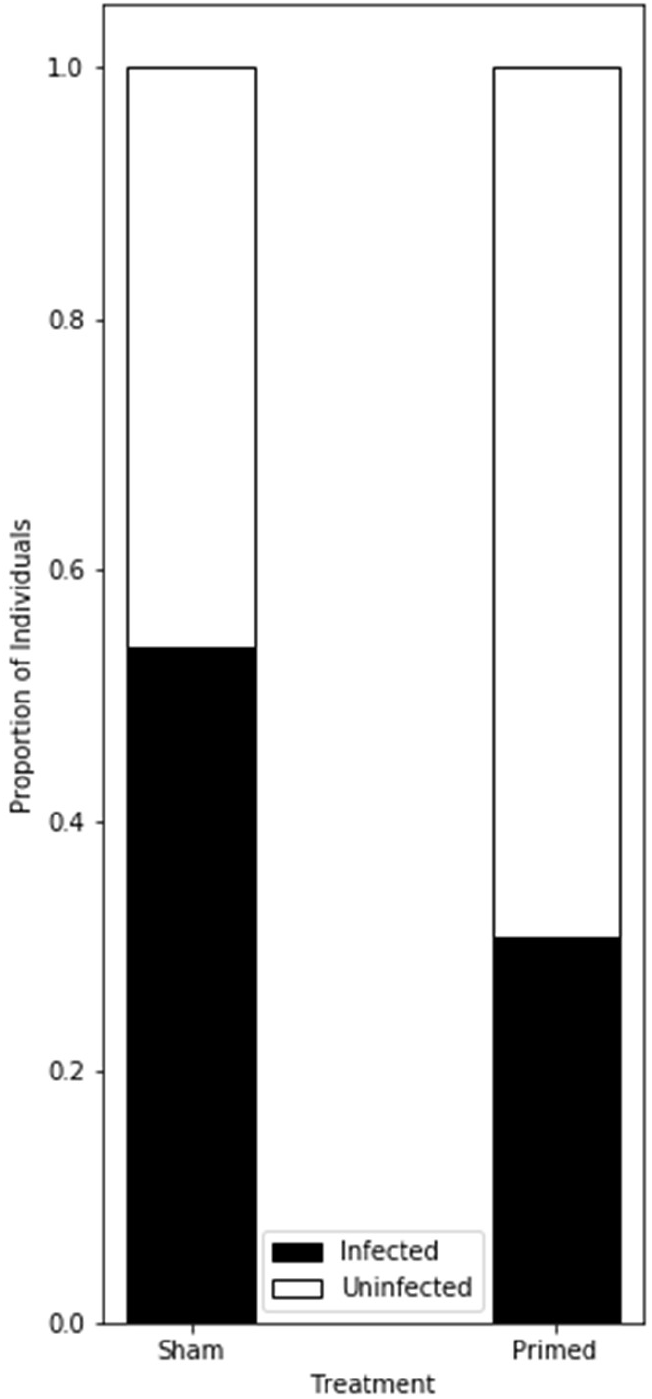
Priming prevents transmissible vectored infection. Approximately half as many Culex quinquefasciatus mosquitoes had detectable West Nile virus in their saliva based on qRT-PCR of the expectorate, (P < 0.05, Fisher’s exact test) when they were given a priming vaccine than those given a sham vaccination. (primed n=78; sham n=76).

### Flavivirus responses

The transcriptomic analyses identified many differentially expressed genes between our baseline control individuals (sham primed–fed control virus free meal) and those given a sham priming and infectious meal (sham–WNV meal, sham–JEV meal). The vast majority of these genes are down-regulated in both WNV meals and JEV meals (Fig 2) and are shared among the flavivirus infections (Fig 3). There are more shared genes that are down-regulated (88.8% [286/322] of WNV and 86.7% [286/330] of JEV) than are up-regulated (5.5% [8/145] WNV, 53.3% [8/15] JEV, Fig 4, Supplemental Table 1). We find that a G-protein coupled receptor gene (CPIJ014334 aka GPROP12, Pteropsin) is among the most strongly suppressed in both the WNV and JEV exposure relative to unexposed mosquitoes. Numerous immune genes are strongly suppressed on WNV infection including Defensin (CPIJ001276, −6.08 log2 fold change (LFC)) Cecropins (CPIJ010699 −5.83 LFC, CPIJ010700, −5.92 LFC), Gram-negative binding protein (CPIJ004324, −5.58 LFC, CPIJ004321, −3.58 LFC), Gambacin (CPIJ016084, −3.46 LFC).

**Fig 2.**
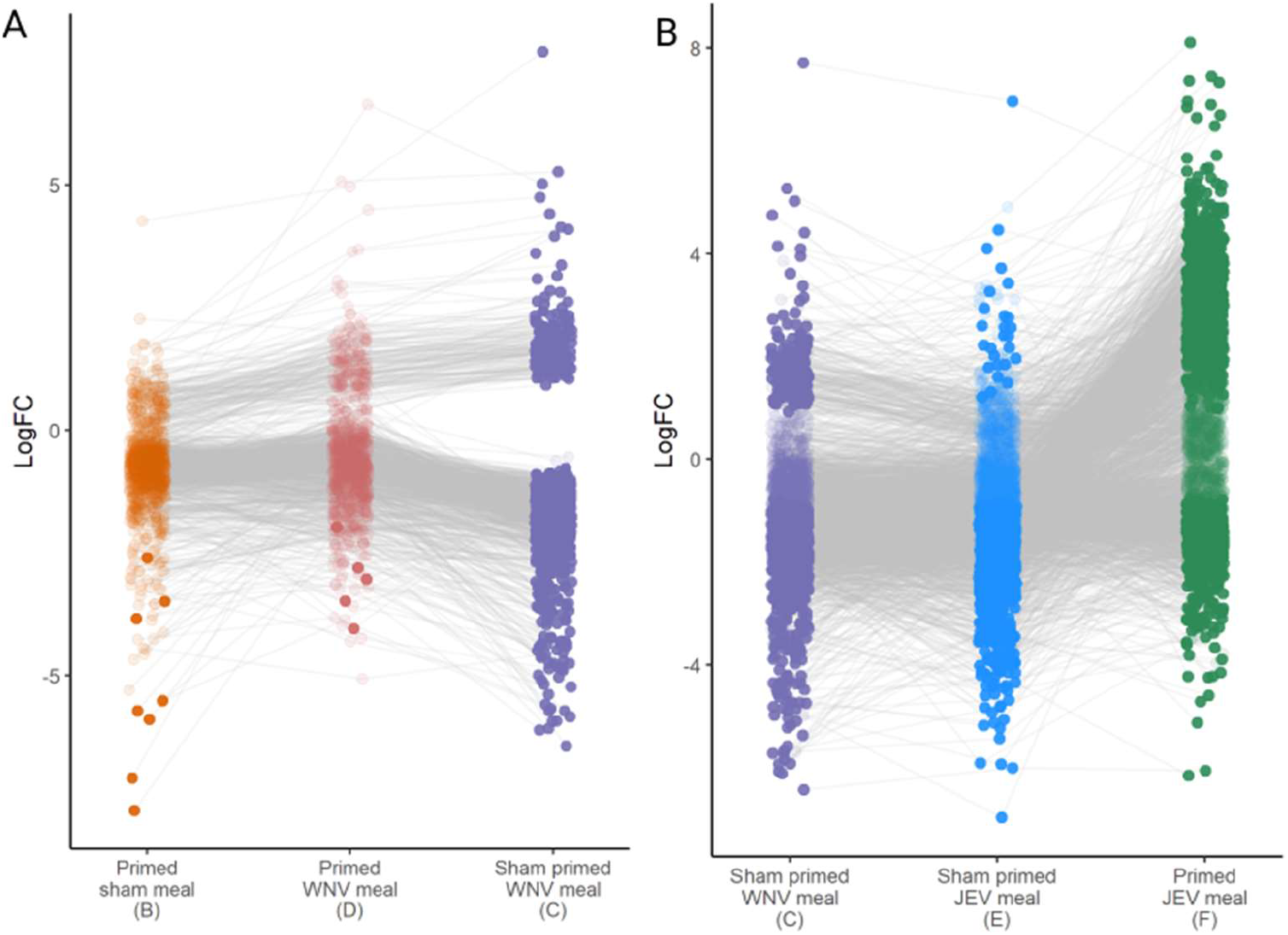
Flavivirus exposure suppresses expression. All expression values (log fold change) are relative to sham-primed individuals given a non-infectious blood meal (condition A). Thus, values below 0 reflect lower expression in these individuals than the control individuals. Figure A) shows expression of mosquitoes primed against WNV and fed virus free meal (B), primed and fed an infectious meal (D) and sham primed and given an infectious meal (C). Figure B. shows the same group C relative to those given a sham priming dose and a JEV infectious meal (E) and those primed against WNV and fed an infectious JEV meal (F). Pale points are not significantly differentially expressed under that condition. Genes connected by grey lines are the same genes in each treatment.

**Fig 3.**
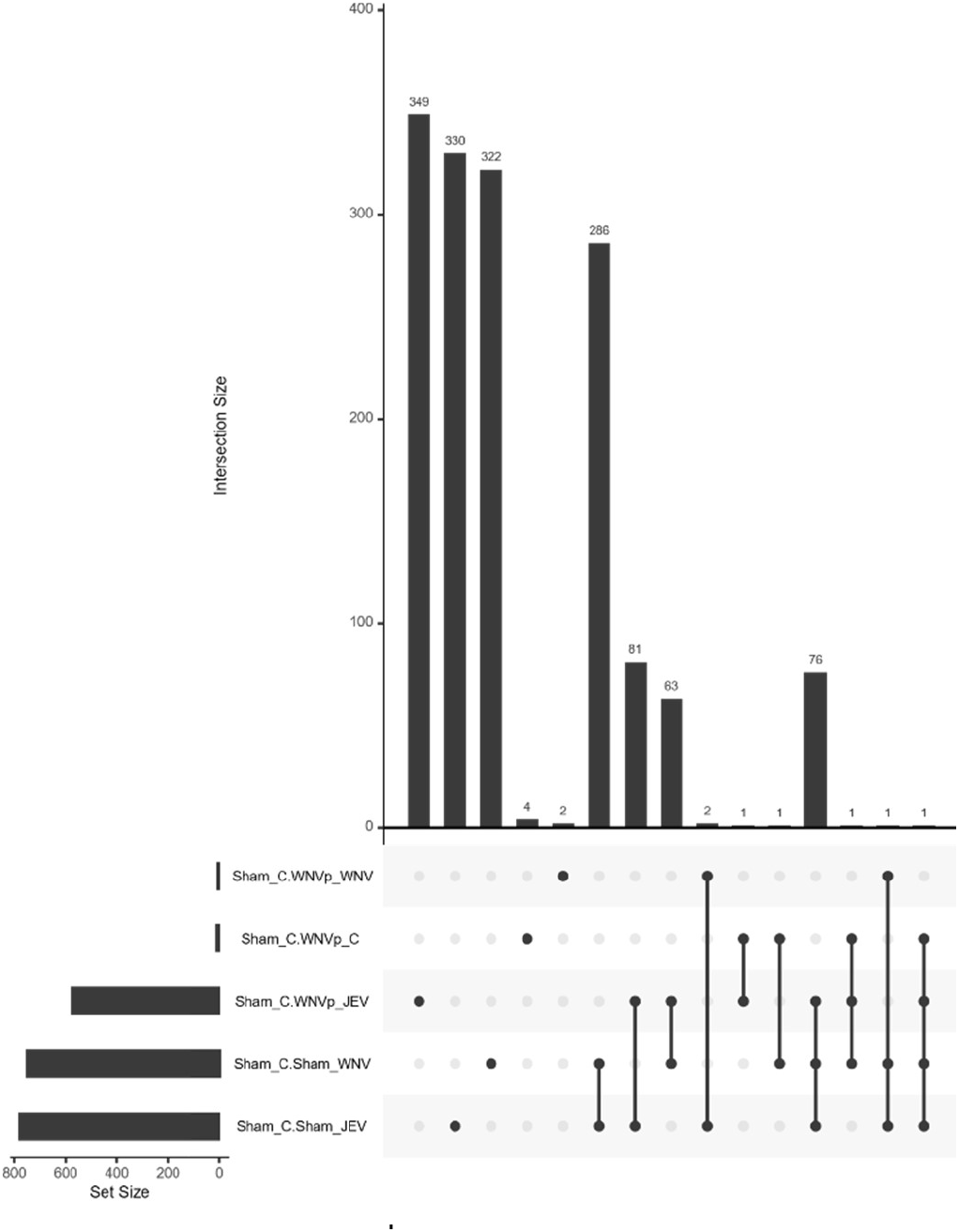
Shared down-regulated genes among flavivirus infected mosquitoes. The size of the bars describe the number of shared genes within that set (designated below the vertical barchart). We only present comparisons (rows) for which there are significant down-regulated genes. An interactive can be accessed at https://bite-and-sting.github.io/mosquitoPriming/

**Fig 4.**
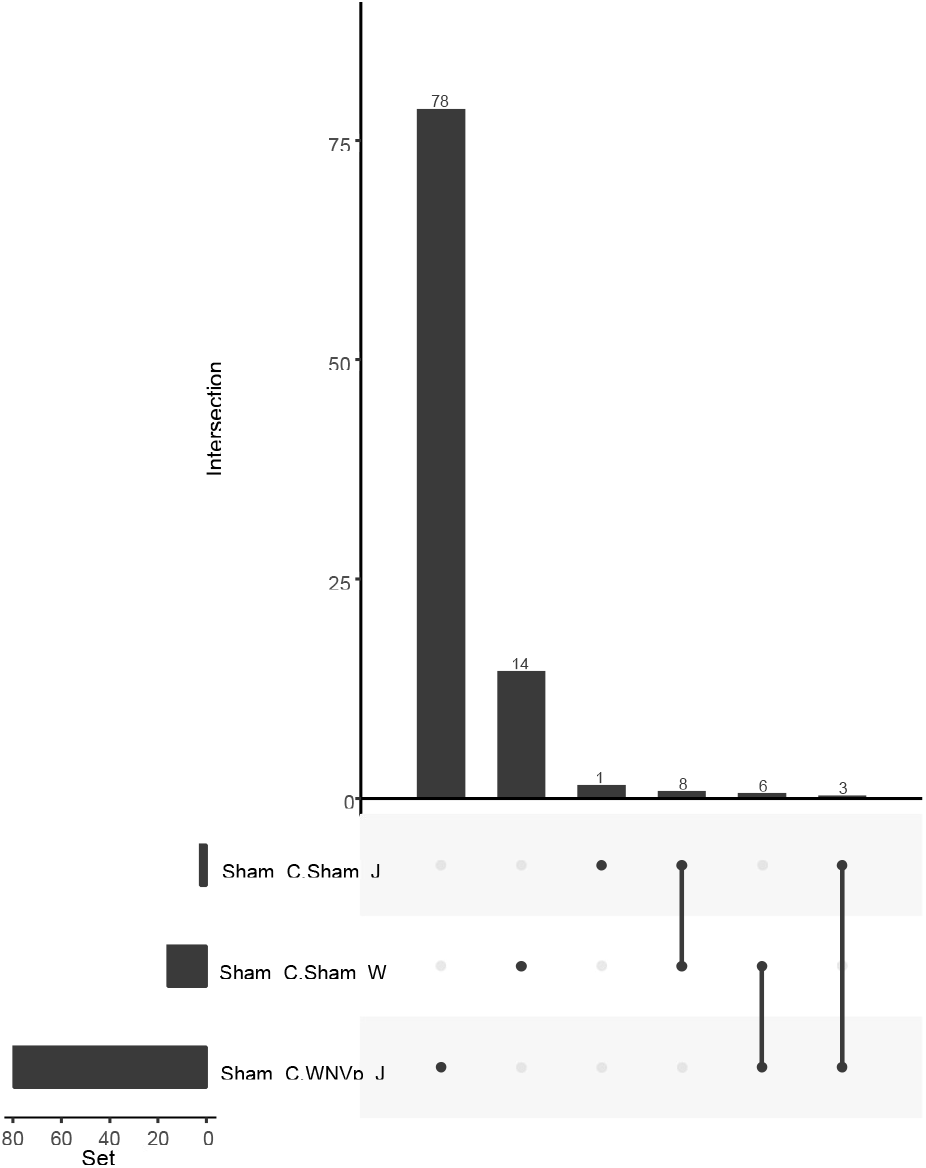
Shared up-regulated genes among flavivirus infected mosquitoes. The size of the bars describe the number of shared genes within that set (designated below the vertical barchart). We only present comparisons (rows) for which there are significant up-regulated genes.

Gene ontology (GO) analyses suggest that infection by WNV results in suppression of a broad suite of genes that encode for functions including immunity, behaviour, sleep, and chemotaxis (Supplemental Table 2). Many, but not all, of these GO terms are shared with JEV exposed mosquitoes. Notably, immune response associated genes are absent in JEV exposed mosquitoes (at *P* < 0.05). A set of these genes are differentially expressed here upon WNV exposure and similarly regulated in the literature (76 differentially expressed genes on WNV exposure in our study are also differentially expressed in our comparison literature data sets, Figure 5, Supplemental Table 3) [37–39] with strong shared over-representation of ontology terms for immunological responses upon WNV exposure, driven by similarity of response to Shin et al.’s [38] and to a lesser extent Bartholomay’s [37] studies. Cecropin is suppressed in both our and Shin et al.’s study, and differentially expressed in Bartholomay et al. although the direction is not given in their supplementary tables.

**Fig 5.**
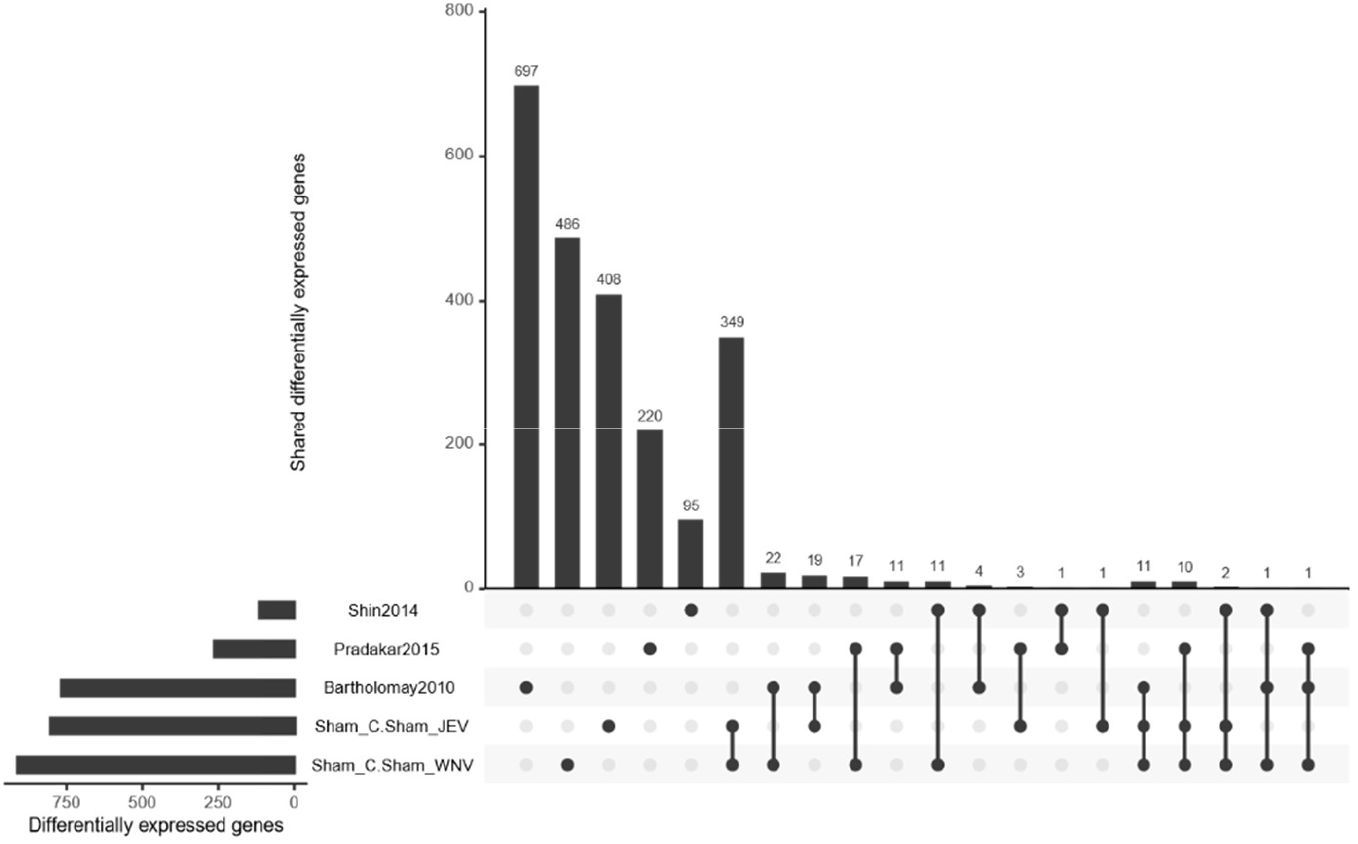
Shared differentially expressed genes on WNV and JEV exposure in this study and those of Bartholomay et al 2010 [18], Paradakar et al 2015 [20], and Shin et al 2014 [19]. The size of the bars describe the number of shared genes within that set (designated below the vertical barchart).

We see further similarity in metabolic and protein localisation and processing in our and Bartholomay’s work. There was overlap in differentially expressed genes in response to JEV exposure from our study and those in the literature (47 genes). The full GO enrichment results are in (Supplemental Table 2) and the overlap between our results and other studies in (Supplemental Table 4). Interactive versions of the overlap plots (Fig 3–5) can be accessed at https://bite-and-sting.github.io/mosquitoPriming/.

### Priming responses

#### Constitutive response

Mosquitoes that received the priming exposure but did not receive an infectious meal (B) had very few differentially expressed genes relative to the sham, unexposed mosquitoes (A) (Fig 6). Of the named genes in this list, we find two immune genes, the lysozyme CPIJ005450 and a Gram-negative binding protein CPIJ004321. All differentially expressed genes under this condition were down-regulated relative to the control group.

**Fig 6.**
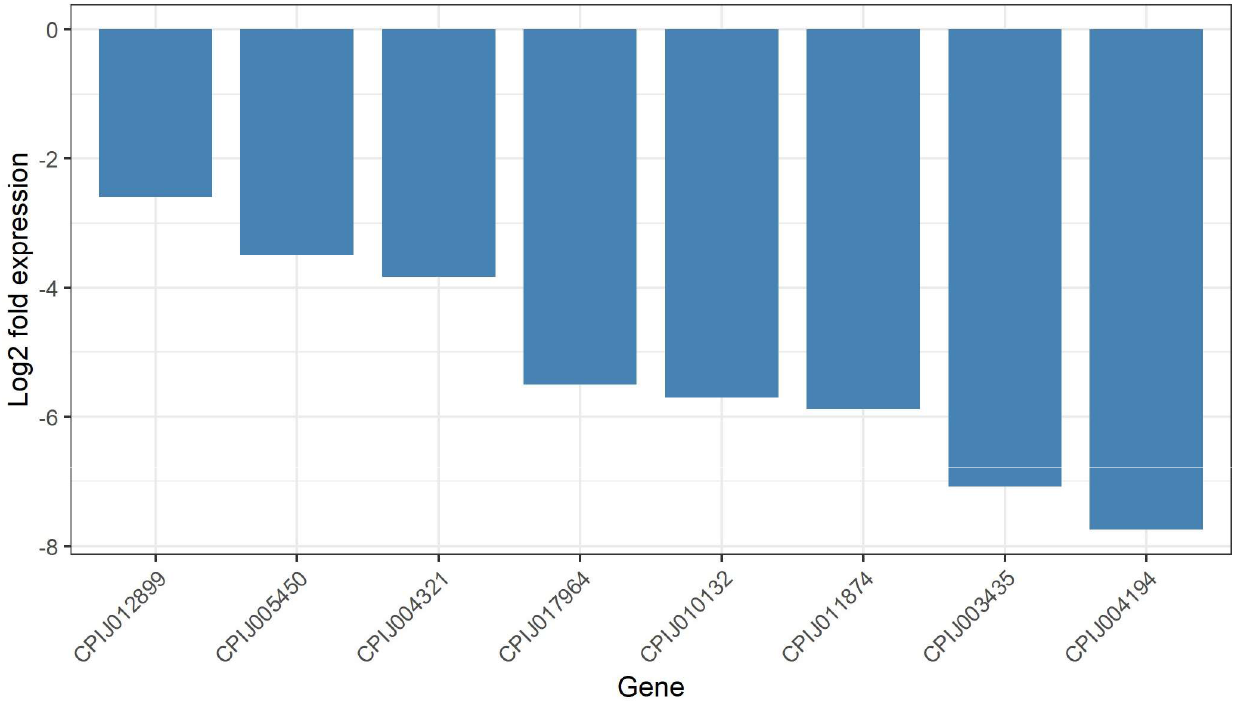
Differentially expressed genes under constitutive memory. Values below 0 reflect lower expression in immune primed but unexposed to live virus than unprimed-unexposed mosquitoes. The genes with annotation include CPIJ005450: Lysozyme, CPIJ004321: Gram-negative bacteria binding protein, CPIJ017964:trypsin 7 precursor, CPIJ003435: Aquarius, CPIJ004194: allergen 2C putative (A vs B).

#### Induced priming responses

Secondary exposure to a WNV infectious meal results in differential expression of 76 genes relative to direct exposure without priming. All but three of which are more highly expressed upon secondary exposure (D) than when mosquitoes are exposed to the infectious meal without the benefit of prior history (C, Fig 7). Among the up-regulated genes are a suite of antimicrobial peptides (CPIJ010700: Cecropin A2, CPIJ010699: Cecropin A1, CPIJ001276: Defensin A). The most strongly upregulated gene on secondary exposure is CPIJ005651 at 7.1 LFC, which equates to greater than 136 fold upregulation on secondary exposure than on primary exposure to WNV. Gene ontology analyses find that as a group, these genes encode responses to infection, detoxification, RNA processes, and stress. The strongest down-regulated gene, gooseberry (CPIJ006496) is a transcription factor which has a number of GO terms that suggest negative regulation of several processes. To determine whether these differences in expression were due to exceptionally high expression on matching priming to exposure conditions, we additionally compared these mosquitoes to the control condition (A vs D) and found very few differentially expressed genes (Suppl. Fig 1) suggesting a return to control-like gene expression.

**Fig 7.**
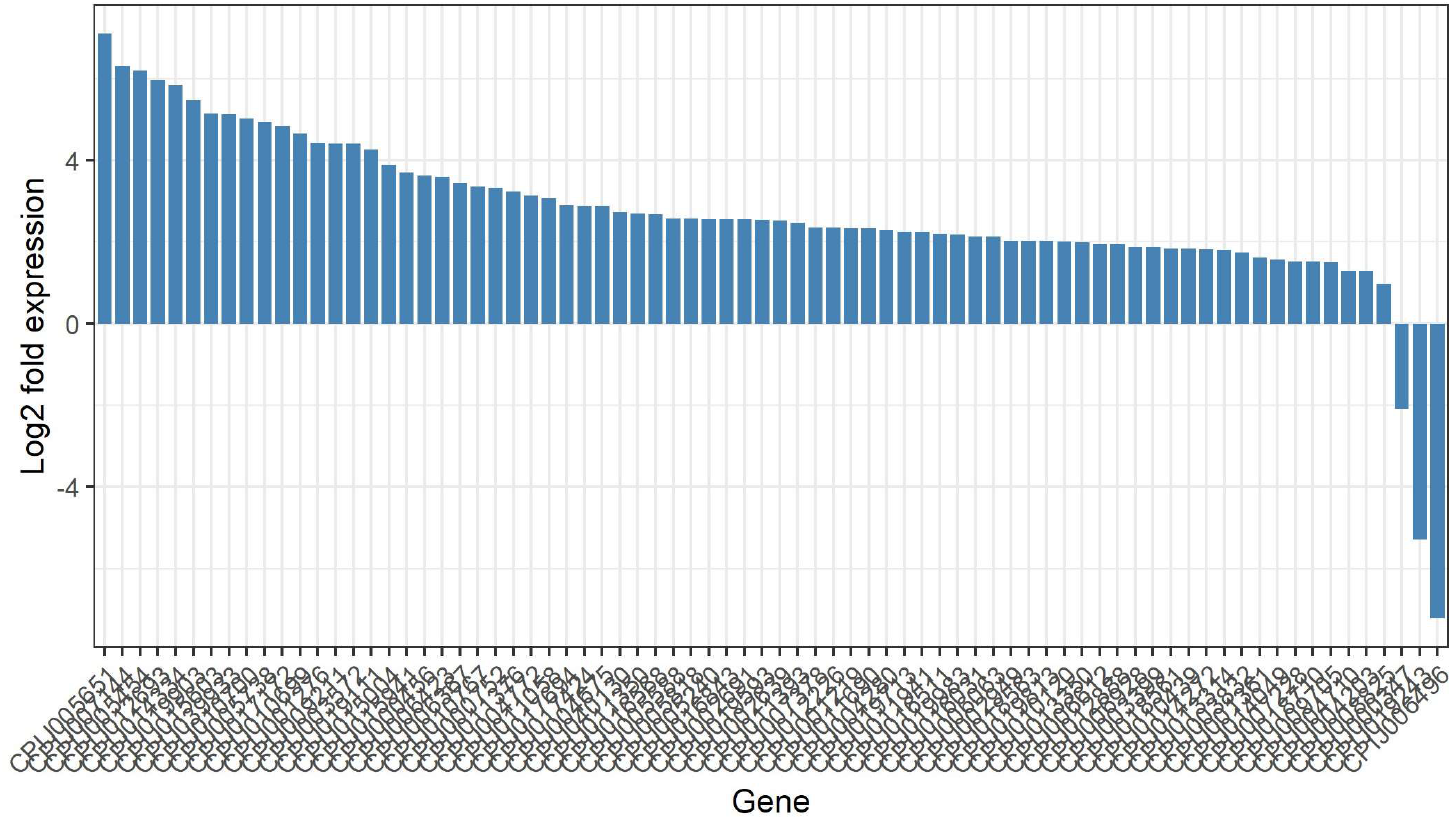
Differentially expressed genes under secondary exposure (exposure matches prior exposure). Values above 0 reflect greater gene expression when mosquitoes have had immune priming and are given an infectious WNV meal than when they have no memory but unexposed to live virus (C vs D).

### Mismatch between priming and exposure

When a mosquito was exposed to JEV but had been primed against WNV it resulted in a vast change in expression with 2,473 differentially expressed genes when comparing the mismatch condition (F) to the normal JEV response (E). This mismatched response was also very different to the matched (D) response with 1,748 differentially expressed genes between groups D and F. The ontology terms associated with these differentially expressed genes were also diverse. In both DF and EF comparisons we find that up-regulated genes on mismatch between priming exposure and infection exposure are involved in RNA processing, RNA metabolism, RNA splicing, and gene expression, immune responses and metabolism. Among the downregulation associated GO terms we find many regulatory terms, protein modification, metabolic processes, and cellular components and cycle elements.

### Co-expression networks

Across all treatments we identify four modules of co-expressed genes of 869, 473, 461, and 74 genes respectively with an additional 12 uncorrelated genes. Broadly, the contribution of these networks can be seen in (Fig 9, Supplemental Table 5). Module 1 consists of genes involved in serine peptidase, cell adhesion, and carbohydrate and lipid metabolism, module 2 includes genes associated with DNA replication, epigenetic processes, chromosomal organization, microtubule movement, and post-transcriptional RNA silencing and includes Maelstrom (CPIJ001566) as a major hub gene. Module 3 includes the antimicrobial peptides (Defensin, Cecopin A1 and A2) and include the small RNA interacting PIWI as a hub gene. Module 4 includes similar terms as those in module 2 (e.g. chromatin, cell cycle, microtubule).

**Fig 8.**
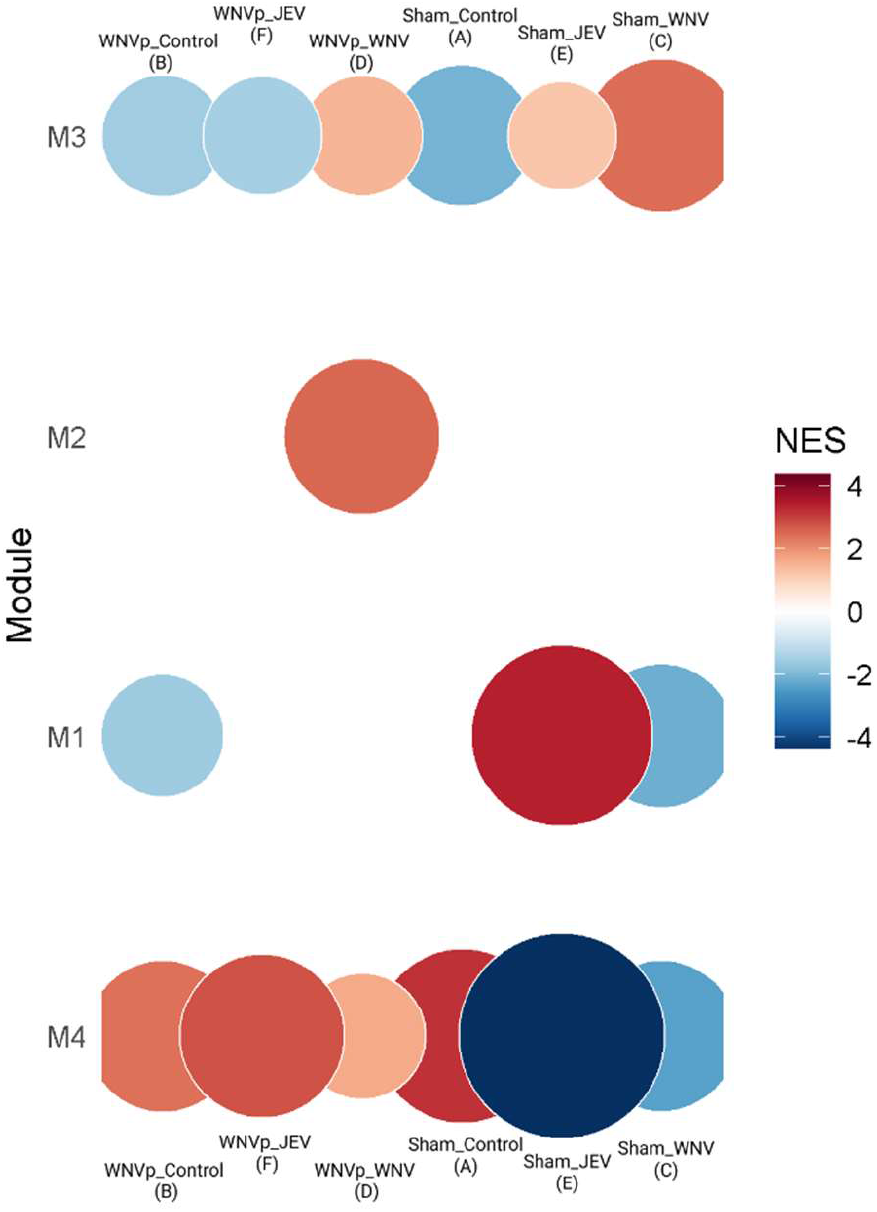
Results of gene set enrichment analysis of members of four dominant modules identified in the CEMiTool network analyses. Names and codes for each treatment are given above and below the plot. The component of the treatment name before the underscore ‘_’ describes the priming treatment (primed with WNV [WNVp] or sham primed). The treatment name after the ‘_’ describes the infection treatment (WNV or JEV blood meals, or blood meal without virus as control). The size of the circle reflects the number of genes

**Fig 9.**
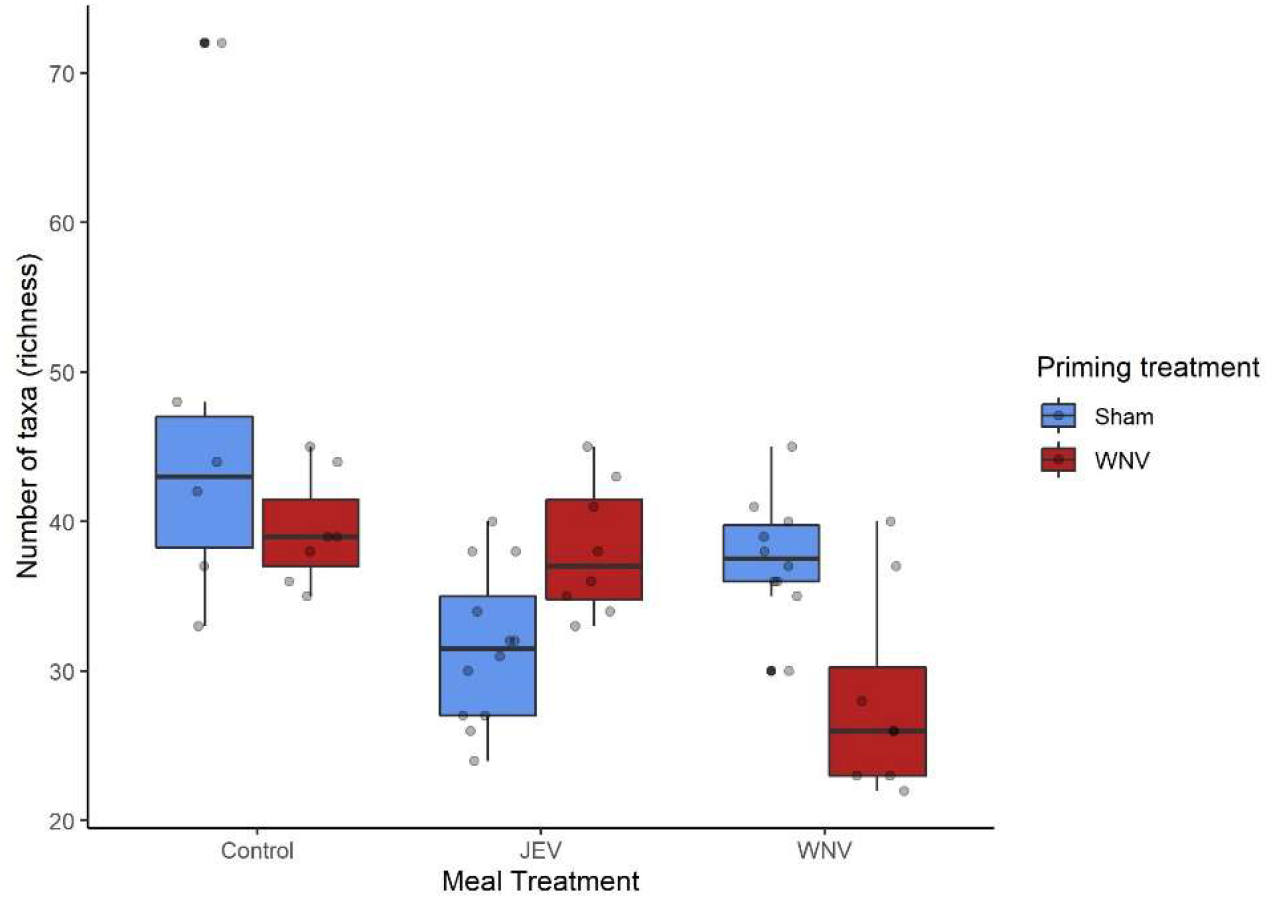
Diversity of non–host reads differs according to treatment. Priming with WNV did not alter the richness of microbial (bacterial and viral) reads that we detect in these samples unless those individuals also receive a live infectious meal of WNV. This result suggests that the induced second response is broad spectrum and alters the community diversity of commensals associated with adult mosquitoes.

Exposure to our two flaviviruses, WNV (C) and JEV (E), resulted in the regulation of fundamentally different networks of genes. WNV infection resulted in stronger up-regulation of genes in module 3 but down-regulation of the genes in modules 1 and 4. JEV saw directionally similar but weaker responses in module 3, and stronger suppression of genes in module 4. Module 1, however was strongly up-regulated in JEV exposure where it was suppressed in WNV exposure. Mosquitoes that were primed and received infectious meal of WNV were unique in their strong regulation of genes in module 2. Other modules show intermediate expression between control (A) and direct exposure (C) groups.

### Microbial diversity

We found that treatments altered the diversity of microbes associated with mosquitoes. While this should be viewed with some caution as we did not set out to describe the complete metagenome of mosquitoes under these conditions, we do find that microbial diversity is lower in mosquitoes that received WNV infectious meals but only when the mosquitoes have had previous exposure to the virus by priming (Fig 9, Supplemental Table 6).

## Discussion

Immune priming can prevent WNV establishment (Fig 1). Here we describe how immune priming alters gene expression with and without receiving a subsequent WNV or JEV infectious meal. We find overlapping but distinct responses to these Flaviviruses. Priming results in increased expression of key effectors including antimicrobial peptides (Defensin, Cecropin A1, Cecropin A2, Diptericin, Gambicin) and genes responsible for epigenetic regulation, post-transcriptional gene silencing, and chromosomal processes. Overall, primed mosquitoes look more like uninfected mosquitoes based on their gene expression. In contrast, Flavivirus infection without immune priming results in wide-scale down-regulation of diverse genes including AMPs and genes responsible for gene regulation.

Here we show that WNV infection broadly suppresses immune gene expression. Of particular note, many AMPs are down-regulated following flavivirus infection. AMPs are active against diverse pathogens including bacteria [42], *Plasmodium* [42], and viruses [43,44]. Here we see possible suppression of AMP expression after WNV infectious meal along with other immunologically important immune effector (lysozyme, ferritin, transferrin, galectin) and recognition genes (PGRP, GNBP) that are linked to activating both Toll and Imd pathways. Pathogens can suppress the immune systems of their hosts, thus ensuring pathogen establishment, proliferation, and ultimately transmission. Pathogen induced immunosuppression has been described in vertebrates[45] and invertebrates [46–48].

In *Cx. quinquefasciatus* cell culture, WNV infection increases Cullin 4 (CPIJ010574) expression andenhances viral proliferation, likely by suppression of Jak-STAT [39]. Here, we find 2.25-fold upregulation of a different Cullin (CPIJ003090) in mosquitoes fed WNV relative to control treated mosquitoes. The *Drosophila melanogaster* ortholog of CPIJ003090 suppresses peptidoglycan recognition and the Imd pathway and AMP production [49]. Strikingly, suppression of Cullin 3 in the plant model species *Arabidopsis thaliana* results in greater basal protection from the bacterial leaf pathogen *Pseudomonas syringae* pv. *maculicola* but incapacitates the systemic acquires response – the plant analog to immune priming [50,51]. We hypothesise a potential mechanism whereby WNV increases Cullin expression, or other immunomodulatory genes, in turn reducing AMP expression. Primed mosquitoes do not succumb to this manipulation, producing stronger immune responses with broad-spectrum effectors such as AMPs. This general response is tentatively supported by our microbial community data (Fig 9) where priming alone does not suppress bacterial and viral diversity, but when mosquitoes are primed and given an infectious dose of WNV the number of detectable taxa dropsdrops to an average of 28.1 taxa from 39.4 in primed but unexposed mosquitoes.

Primed mosquitoes appear to be able to avoid such putative viral manipulation and this may be evident in our network analysis, where infection up-regulates genes in both module 2 and 4 which constitutes a swathe of genes with regulatory functions including post-transcriptional RNA silencing, chromatin, histone modification, DNA replication and chromosome organisation. Previous work has described the role of DNA replication in producing primed immune responses of *Anopheles albimanus* cell lines to *Plasmodium berghei* [52], and *Aedes aegypti* to Dengue infection [15,53,54]. Our finding, using whole transcriptome sequencing, finds increased expression of a Delta-like gene on WNV exposure (CPIJ019430), which may be the ortholog of a gene implicated in priming by the studies of Serrato–Salas [53,54], but we did not find altered expression of Notch or Hindsight which they also link to priming in *Ae. aegypti*. Nor do we find differential expression of these genes in any of our primed conditions (B,D,F). Expanding this survey to all genes in the KEGG pathway for *Cx. quinquefasciatus* Notch signalling, we find none of these genes are differentially expressed in any of our comparisons.

The hypothetical protein CPIJ005651 is strongly upregulated on exposure to WNV in primed mosquitoes (D). This gene is not significantly suppressed in WNV infected, unprimed, mosquitoes (C), in contrast to the AMPs described above. CPIJ005651 is an ortholog of the *D. melanogaster* cap-binding protein 80 (Cbp80, Fbgn002942, OrthoDB ID: 16059at6960); a highly conserved protein that regulates mRNA and miRNA production, RNA interference (RNAi), and nonsense mediated RNA decay. There appears to also be a role for Cbp80 and other members of the cap binding protein complex, and nonsense-mediated decay process more generally, in viral immunity in yeast [55], plants [56], fruit flies [57], mice [58,59]. Cbp80 is required for small interfering RNA response to several *D. melanogaster* RNA viruses [57]. RNAi based knock down of Cbp80 in *D. melanogaster* cell culture resulted in higher titers of several RNA viruses including *Drosophila* C-virus and vesicular stomatitis virus (VSV) and comparable manipulation of the protein interacting partner, Ars2 results in similarly increased infection in adult flies.

The role of Cap binding proteins in viral defense seems to be ancient as orthologs, or members of the same complex, have been implicated in defense against plant viruses [56], and in vertebrate model systems [58,59]. Cap binding proteins and Ars2 have not yet been described as important for mosquito viral responses but our finding that CPIJ005651 is strongly upregulated during priming raises the potential that it plays a role in viral immunity, as it does in evolutionarily divergent systems. Cbp80 and associated genes may contribute to both altering miRNA and mRNA abundances that confer greater protection against secondary WNV infection, or could hypothetically directly suppress viral replication in host cells by degrading capped viral RNA. Beyond putative direct anti-viral responses, cap binding processes may further play a role in immune priming. It is striking that EIF4-e domain containing gene was the most strongly differentially expressed gene in trans-generationally primed *Tribolium castaneum* [60], and is the major target of mTOR in the mammalian trained innate immune memory of macrophages [61]. In our study, we find that expression of EIF4-e (CPIJ012031) is suppressed on both WNV and JEV infection but returns to control levels when mosquitoes had been primed and unexposed (B) or receive matching exposure (D).

In summary, we find that flavivirus infection suppresses vector gene expression, including key immune genes such as AMPs, but that prior immune exposure to WNV can rescue expression from viral suppression. This evasion of viral suppression, plausibly through RNA metabolic and transfer mechanisms, could explain the reduced infection of primed mosquitoes in our experiment. While these findings raise exciting potential to understand immunity and how it can be applied to control of vector bourn infections, much still needs to be clarified. Among the questions to be addressed include the importance of dose and delivery to the degree of protection. Immune priming can be produced by consumed, injected, and even inherited exposures. The relative protection of these different exposures is important to clarify. The crucial question that remains unanswered in vector priming, is how specific is the priming. Here lies the greatest possible opportunity, and the greatest possible risk. Priming can be highly specific [16], and/or can trade-off against protection to other pathogens [60] and other life history traits [62]. Both specificity and the trade-offs are relevant and important to understand in vectors. On one hand, priming – if general – could help protect against multiple vectored diseases; on the other – if resistances trade-off – priming could increase susceptibility to other pathogens. The greater susceptibility to other infections may include vectored disease but also ecologically relevant entomopathogens.

We acknowledge certain limitations to our study as follows:

1. Time post priming. We challenged mosquitoes with an infectious blood meal at seven-days post priming. This time point was chosen as the initial immune response to the physical damage would have subsided [63], and also to give mosquitoes time to recover following injection enabling them to feed properly and mount an immunologically ‘informed’ response. Indeed, we see few genes differentially expressed after priming alone. It is feasible that the response to priming varies over time, while shorter time-points may lead to artefacts, testing the effect in older (and/or repeatedly fed) mosquitoes should be investigated following on from our proof-of-principle presented here.
2. Dose and delivery effects. Here we examined the priming protection and gene expression of mosquitoes given a single priming dose and exposed to a single infectious meal at one dose of exposure. The intensity of priming and degree of protection may well vary according to the amount of virus in the priming exposure and may also be attainable with oral priming, which is a promising direction for subsequent study.
3. Single mosquito species. We have only demonstrated the effect of priming in a single species of mosquito, *Cx. quinquefaciatus*. Both JEV and WNV have other vectors for which the effect of priming may differ. However, *Cx. quinquefaciatus* is an important vector of both viruses, in both of their reservoir cycles and spill-over to humans[4–6]. Future work should identify the effect of priming in other species, such as *Cx. pipiens*, and *Cx. tarsalis* for WNV. Both of these species are significantly more difficult to manipulate in the laboratory (especially *Cx. pipiens*), and genomic resources less well developed.
4. While RNA sequencing does provide insight into the genes that respond to infection and priming, further functional assays (e.g. knockdown) would be necessary to conclusively determine responsible agents of immune priming.

## Conclusion

WNV infection results in wide-scale suppression of gene expression including crucial antimicrobial peptides but prior exposure to inactivated WNV prevents subsequent infection. Thus, while no human vaccine to WNV exists, we have demonstrated the potential for vaccinating vectors against this virus. We show that immune priming appears to prevent this viral manipulation of host immune expression, identifying a plausible route to priming-induced resistance. The two Flaviviruses tested produce distinct expression profiles in their mosquito vectors with comparably distinct co-expression networks. Our study describes, for the first time, the transcriptomic responses of mosquitoes to viral priming and subsequent exposure to Flavivirus infection. Priming can effectively suppress vector infection to several diseases of human importance although the full diversity remains unknown. Further work to diversify the taxonomic breadth of vectored diseases that priming can prevent and a means to efficiently implement priming in the field stands to add a valuable tool to the control of vector-borne infectious diseases of human and non-human animals.

## Supporting information

Supplements can be found in github: https://github.com/Bite-and-Sting/mosquitoPriming

**S1 Table. Differentially expressed genes in each comparison.**

**S2 Table. Overrepresented gene ontology (GO) terms in each comparison.**

**S3 Table. Overlap in differentially expressed genes in this study, and those of other published studies** [37–39].

**S4 Table. Overlap in overrepresented GO terms found in this study and those of other published studies** [37–39].

**S5 Table. Network tables that describe co-expressed genes.**

**S6 Table. Non-eukaryote taxa found in mosquitoes based on sequence reads.**

## Acknowledgments

This work was supported by Wellcome Trust grant (212450/Z/18/Z) to SMB and MB. We thank Ben M Sadd and Jandouwe Villinger for feedback on an earlier version of this manuscript.

## References

1. WHO. Burden of Vector Borne Diseases Annex I. 2017. Available: https://www.who.int/vector-control/burden_vector-borne_diseases.pdf?ua=1

2. Blagrove MSC, Caminade C, Diggle PJ, Patterson EI, Sherlock K, Chapman GE, et al. Potential for Zika virus transmission by mosquitoes in temperate climates. Proc Biol Sci. 2020;287: 20200119.

3. Gubler DJ. The continuing spread of West Nile virus in the western hemisphere. Clin Infect Dis. 2007;45: 1039–1046.

4. Ciota AT. West Nile virus and its vectors. Curr Opin Insect Sci. 2017;22: 28–36.

5. Pearce JC, Learoyd TP, Langendorf BJ, Logan JG. Japanese encephalitis: the vectors, ecology and potential for expansion. J Travel Med. 2018;25: S16–S26.

6. Garcia-Rejon JE, Blitvich BJ, Farfan-Ale JA, Loroño-Pino MA, Chi Chim WA, Flores-Flores LF, et al. Host-feeding preference of the mosquito, Culex quinquefasciatus, in Yucatan State, Mexico. J Insect Sci. 2010;10: 32.

7. Gould E, Pettersson J, Higgs S, Charrel R, de Lamballerie X. Emerging arboviruses: Why today? One Health. 2017;4: 1–13.

8. Blagrove MSC, Arias-Goeta C, Failloux A-B, Sinkins SP. Wolbachia strain wMel induces cytoplasmic incompatibility and blocks dengue transmission in Aedes albopictus. Proc Natl Acad Sci U S A. 2012;109: 255–260.

9. Walker T, Johnson PH, Moreira LA, Iturbe-Ormaetxe I, Frentiu FD, McMeniman CJ, et al. The wMel Wolbachia strain blocks dengue and invades caged Aedes aegypti populations. Nature. 2011;476: 450–453.

10. Williams AE, Franz AWE, Reid WR, Olson KE. Antiviral Effectors and Gene Drive Strategies for Mosquito Population Suppression or Replacement to Mitigate Arbovirus Transmission by Aedes aegypti. Insects. 2020;11. doi:10.3390/insects11010052

11. Milutinović B, Kurtz J. Immune memory in invertebrates. Semin Immunol. 2016. doi:10.1016/j.smim.2016.05.004

12. Roth O, Beemelmanns A, Barribeau SM, Sadd BM. Recent advances in vertebrate and invertebrate transgenerational immunity in the light of ecology and evolution. Heredity. 2018;121: 225–238.

13. Rodrigues J, Brayner FA, Alves LC, Dixit R, Barillas-Mury C. Hemocyte differentiation mediates innate immune memory in Anopheles gambiae mosquitoes. Science. 2010;329: 1353–1355.

14. Ramirez JL, de Almeida Oliveira G, Calvo E, Dalli J, Colas RA, Serhan CN, et al. A mosquito lipoxin/lipocalin complex mediates innate immune priming in Anopheles gambiae. Nat Commun. 2015;6: 7403.

15. Vargas V, Cime-Castillo J, Lanz-Mendoza H. Immune priming with inactive dengue virus during the larval stage of Aedes aegypti protects against the infection in adult mosquitoes. Sci Rep. 2020;10: 6723.

16. Roth O, Sadd BM, Schmid-Hempel P, Kurtz J. Strain-specific priming of resistance in the red flour beetle, Tribolium castaneum. Proc Biol Sci. 2009;276: 145–151.

17. Sadd BM, Schmid-Hempel P. A distinct infection cost associated with trans-generational priming of antibacterial immunity in bumble-bees. Biol Lett. 2009;5: 798–801.

18. van den Hurk AF, Ritchie SA, Mackenzie JS. Ecology and geographical expansion of Japanese encephalitis virus. Annu Rev Entomol. 2009;54: 17–35.

19. Japanese encephalitis. [cited 5 Nov 2020]. Available: https://www.who.int/news-room/fact-sheets/detail/japanese-encephalitis

20. Campbell GL, Hills SL, Fischer M, Jacobson JA, Hoke CH, Hombach JM, et al. Estimated global incidence of Japanese encephalitis: a systematic review. Bull World Health Organ. 2011;89: 766–74, 774A-774E.

21. Tsai T. New initiatives for the control of Japanese encephalitis by vaccination:: Minutes of a WHO/CVI meeting, Bangkok, Thailand, 13–15 October 1998. Vaccine. 2000;Supplement 2: 1–25.

22. Solomon T, Ooi MH, Beasley DWC, Mallewa M. West Nile encephalitis. BMJ. 2003;326: 865–869.

23. Chancey C, Grinev A, Volkova E, Rios M. The global ecology and epidemiology of West Nile virus. Biomed Res Int. 2015;2015: 376230.

24. Weiss D, Carr D, Kellachan J, Tan C, Phillips M, Bresnitz E, et al. Clinical findings of West Nile virus infection in hospitalized patients, New York and New Jersey, 2000. Emerg Infect Dis. 2001;7: 654–658.

25. Fang Y, Brault AC, Reisen WK. Comparative thermostability of West Nile, St. Louis encephalitis, and western equine encephalomyelitis viruses during heat inactivation for serologic diagnostics. Am J Trop Med Hyg. 2009;80: 862–863.

26. Hadfield TL, Turell M, Dempsey MP, David J, Park EJ. Detection of West Nile virus in mosquitoes by RT-PCR. Mol Cell Probes. 2001;15: 147–150.

27. Dobin A, Davis CA, Schlesinger F, Drenkow J, Zaleski C, Jha S, et al. STAR: ultrafast universal RNA-seq aligner. Bioinformatics. 2013;29: 15–21.

28. McCarthy DJ, Chen Y, Smyth GK. Differential expression analysis of multifactor RNA-Seq experiments with respect to biological variation. Nucleic Acids Res. 2012;40: 4288–4297.

29. Robinson MD, McCarthy DJ, Smyth GK. edgeR: a Bioconductor package for differential expression analysis of digital gene expression data. Bioinformatics. 2010;26: 139–140.

30. Alexa A, Rahnenfuhrer J. stopGO: Enrichment Analysis for Gene Ontology. 2020. Available: https://www.bioconductor.org/packages/release/bioc/vignettes/topGO/inst/doc/topGO.pdf

31. Russo PST, Ferreira GR, Cardozo LE, Bürger MC, Arias-Carrasco R, Maruyama SR, et al. CEMiTool: a Bioconductor package for performing comprehensive modular co-expression analyses. BMC Bioinformatics. 2018;19: 56.

32. Cheng CW, Beech DJ, Wheatcroft SB. Advantages of CEMiTool for gene co-expression analysis of RNA-seq data. Comput Biol Med. 2020;125: 103975.

33. Szklarczyk D, Gable AL, Lyon D, Junge A, Wyder S, Huerta-Cepas J, et al. STRING v11: protein-protein association networks with increased coverage, supporting functional discovery in genome-wide experimental datasets. Nucleic Acids Res. 2019;47: D607–D613.

34. Raudvere U, Kolberg L, Kuzmin I, Arak T, Adler P, Peterson H, et al. g:Profiler: a web server for functional enrichment analysis and conversions of gene lists (2019 update). Nucleic Acids Res. 2019;47: W191–W198.

35. Gehlenborg N. A More Scalable Alternative to Venn and Euler Diagrams for Visualizing Intersecting Sets [R package UpSetR version 1.4.0]. 2019 [cited 29 Sep 2020]. Available: https://CRAN.R-project.org/package=UpSetR

36. R Development Core Team. R: A Language and Environment for Statistical Computing. Vienna, Austria; 2012. Available: http://www.R-project.org

37. Bartholomay LC, Waterhouse RM, Mayhew GF, Campbell CL, Michel K, Zou Z, et al. Pathogenomics of Culex quinquefasciatus and meta-analysis of infection responses to diverse pathogens. Science. 2010;330: 88–90.

38. Shin D, Civana A, Acevedo C, Smartt CT. Transcriptomics of differential vector competence: West Nile virus infection in two populations of Culex pipiens quinquefasciatus linked to ovary development. BMC Genomics. 2014;15: 513.

39. Paradkar PN, Duchemin J-B, Rodriguez-Andres J, Trinidad L, Walker PJ. Cullin4 Is Pro-Viral during West Nile Virus Infection of Culex Mosquitoes. PLoS Pathog. 2015;11: e1005143.

40. Saha S, Ramesh A, Kalantar K, Malaker R, Hasanuzzaman M, Khan LM, et al. Unbiased metagenomic sequencing for pediatric meningitis in Bangladesh reveals neuroinvasive Chikungunya virus outbreak and other unrealized pathogens. bioRxiv. 2019. p. 579532. doi:10.1101/579532

41. Oksanen J, Blanchet FG, Friendly M, Kindt R, Legendre P, McGlinn D, et al. vegan: Community Ecology Package. 2019. Available: https://CRAN.R-project.org/package=vegan

42. Kokoza V, Ahmed A, Woon Shin S, Okafor N, Zou Z, Raikhel AS. Blocking of Plasmodium transmission by cooperative action of Cecropin A and Defensin A in transgenic Aedes aegypti mosquitoes. Proc Natl Acad Sci U S A. 2010;107: 8111–8116.

43. Zhao L, Alto BW, Smartt CT, Shin D. Transcription Profiling for Defensins of Aedes aegypti (Diptera: Culicidae) During Development and in Response to Infection With Chikungunya and Zika Viruses. J Med Entomol. 2018;55: 78–89.

44. Pan X, Zhou G, Wu J, Bian G, Lu P, Raikhel AS, et al. Wolbachia induces reactive oxygen species (ROS)-dependent activation of the Toll pathway to control dengue virus in the mosquito Aedes aegypti. Proc Natl Acad Sci U S A. 2012;109: E23–31.

45. Janeway CA Jr, Travers P, Walport M, Shlomchik MJ. Immunobiology. Garland Science; 2001.

46. Apidianakis Y, Mindrinos MN, Xiao W, Lau GW, Baldini RL, Davis RW, et al. Profiling early infection responses: Pseudomonas aeruginosa eludes host defenses by suppressing antimicrobial peptide gene expression. Proc Natl Acad Sci U S A. 2005;102: 2573–2578.

47. Barribeau SM, Sadd BM, du Plessis L, Schmid-Hempel P. Gene expression differences underlying genotype-by-genotype specificity in a host-parasite system. Proc Natl Acad Sci U S A. 2014;111: 3496–3501.

48. Lefèvre T, Adamo SA, Biron DG, Missé D, Hughes D, Thomas F. Invasion of the body snatchers: the diversity and evolution of manipulative strategies in host–parasite interactions. Adv Parasitol. 2009;68: 45–83.

49. Khush RS, Cornwell WD, Uram JN, Lemaitre B. A ubiquitin-proteasome pathway represses the Drosophila immune deficiency signaling cascade. Curr Biol. 2002;12: 1728–1737.

50. Spoel SH, Mou Z, Tada Y, Spivey NW, Genschik P, Dong X. Proteasome-mediated turnover of the transcription coactivator NPR1 plays dual roles in regulating plant immunity. Cell. 2009;137: 860–872.

51. Fu ZQ, Yan S, Saleh A, Wang W, Ruble J, Oka N, et al. NPR3 and NPR4 are receptors for the immune signal salicylic acid in plants. Nature. 2012;486: 228–232.

52. Cime-Castillo J, Arts RJW, Vargas-Ponce de León V, Moreno-Torres R, Hernández-Martínez S, Recio-Totoro B, et al. DNA Synthesis Is Activated in Mosquitoes and Human Monocytes During the Induction of Innate Immune Memory. Front Immunol. 2018;9: 2834.

53. Serrato-Salas J, Izquierdo-Sánchez J, Argüello M, Conde R, Alvarado-Delgado A, Lanz-Mendoza H. Aedes aegypti antiviral adaptive response against DENV-2. Dev Comp Immunol. 2018;84: 28–36.

54. Serrato-Salas J, Hernández-Martínez S, Martínez-Barnetche J, Condé R, Alvarado-Delgado A, Zumaya-Estrada F, et al. De Novo DNA Synthesis in Aedes aegypti Midgut Cells as a Complementary Strategy to Limit Dengue Viral Replication. Front Microbiol. 2018;9: 801.

55. Kushner DB, Lindenbach BD, Grdzelishvili VZ, Noueiry AO, Paul SM, Ahlquist P. Systematic, genomewide identification of host genes affecting replication of a positive-strand RNA virus. Proc Natl Acad Sci U S A. 2003;100: 15764–15769.

56. Pasin F, Shan H, García B, Müller M, San León D, Ludman M, et al. Abscisic Acid Connects Phytohormone Signaling with RNA Metabolic Pathways and Promotes an Antiviral Response that Is Evaded by a Self-Controlled RNA Virus. Plant Commun. 2020;1. doi:10.1016/j.xplc.2020.100099

57. Sabin LR, Zhou R, Gruber JJ, Lukinova N, Bambina S, Berman A, et al. Ars2 regulates both miRNA- and siRNA-dependent silencing and suppresses RNA virus infection in Drosophila. Cell. 2009;138: 340–351.

58. Gebhardt A, Bergant V, Schnepf D, Moser M, Meiler A, Togbe D, et al. The alternative cap-binding complex is required for antiviral defense in vivo. PLoS Pathog. 2019;15: e1008155.

59. Gebhardt A, Habjan M, Benda C, Meiler A, Haas DA, Hein MY, et al. mRNA export through an additional cap-binding complex consisting of NCBP1 and NCBP3. Nat Commun. 2015;6: 8192.

60. Tate AT, Andolfatto P, Demuth JP, Graham AL. The within-host dynamics of infection in trans-generationally primed flour beetles. Mol Ecol. 2017;26: 3794–3807.

61. Cheng S-C, Quintin J, Cramer RA, Shepardson KM, Saeed S, Kumar V, et al. mTOR- and HIF-1α-mediated aerobic glycolysis as metabolic basis for trained immunity. Science. 2014;345: 1250684.

62. Contreras-Garduño J, Rodríguez MC, Rodríguez MH, Alvarado-Delgado A, Lanz-Mendoza H. Cost of immune priming within generations: trade-off between infection and reproduction. Microbes Infect. 2014;16: 261–267.

63. Kambris Z, Blagborough AM, Pinto SB, Blagrove MSC, Godfray HCJ, Sinden RE, et al. Wolbachia stimulates immune gene expression and inhibits plasmodium development in Anopheles gambiae. PLoS Pathog. 2010;6: e1001143.

